# Compartment-specific energy requirements of photosynthetic carbon metabolism in *Camelina sativa* leaves

**DOI:** 10.1101/2022.01.28.478172

**Authors:** Thomas Wieloch, Thomas David Sharkey

**Affiliations:** Department of Medical Biochemistry and Biophysics, KB. H6, Umeå University, 901 87 Umeå, Sweden; Department of Energy-Plant Research Laboratory, Michigan State University, East Lansing, MI 48824, USA; Department of Biochemistry and Molecular Biology, Michigan State University, East Lansing, MI 48824, USA; Plant Resilience Institute, Michigan State University, East Lansing, MI 48824, USA

**Keywords:** ATP/NADPH ratio, bioenergetics, energy metabolism, glucose-6-phosphate shunt, oxidative pentose phosphate pathway, sucrose cycling

## Abstract

Detailed knowledge about plant energy metabolism may aid crop improvements. Using published estimates of flux through central carbon metabolism, we phenotype energy metabolism in illuminated *Camelina sativa* leaves (grown at 22 °C, 500 μmol photons m^-2^ s^-1^) and report several findings. First, the oxidative pentose phosphate pathway (OPPP) transfers 3.3% of the NADPH consumed in the Calvin-Benson cycle to the cytosol. NADPH supply proceeds at about 10% of the rate of net carbon assimilation. However, concomitantly respired CO_2_ accounts for 4.8% of total rubisco activity. Hence, 4.8% of the flux through the Calvin-Benson cycle and photorespiration is spent on supplying cytosolic NADPH, a significant investment. Associated energy requirements exceed the energy output of the OPPP. Thus, autotrophic carbon metabolism is not simply optimised for flux into carbon sinks but sacrifices carbon and energy use efficiency to support cytosolic energy metabolism. To reduce these costs, we suggest bioengineering plants with a repressed cytosolic OPPP, and an inserted cytosolic NADPH-dependent malate dehydrogenase tuned to compensate for the loss in OPPP activity (if required). Second, sucrose cycling is a minor investment in overall leaf energy metabolism but a significant investment in cytosolic energy metabolism. Third, leaf energy balancing strictly requires oxidative phosphorylation, cofactor export from chloroplasts, and peroxisomal NADH import. Fourth, mitochondria are energetically self-sufficient. Fifth, carbon metabolism has an ATP/NADPH demand ratio of 1.52 which is met if ≤21.7% of whole electron flux is cyclic. Sixth, electron transport has a photon use efficiency of ≥62%. Lastly, we discuss interactions between the OPPP and the cytosolic oxidation-reduction cycle in supplying leaf cytosolic NADPH.

**Main Conclusion:** The oxidative pentose phosphate pathway provides cytosolic NADPH yet reduces carbon and energy use efficiency. Repressing this pathway and introducing cytosolic NADPH-dependent malate dehydrogenase may increase crop yields by ≈5%.

## Introduction

Central carbon metabolism operates co-ordinately across several cell compartments and requires compartment-specific inputs of reductant and energy cofactors (NAD(P)H, Fd_red_, ATP, UTP). These requirements vary with ambient and developmental conditions (Scheibe 2019). For instance, drought frequently causes stomatal closure and low intercellular CO_2_ concentrations which promote photorespiratory NADH production in mitochondria and consumption in peroxisomes (see below). Since cofactor production is spatially restricted, transmembrane transport and conversion of cofactors are key to metabolic functioning.

In illuminated leaves, reductant and energy cofactors are primarily produced by the light reactions of photosynthesis inside chloroplasts. Additionally, the pyruvate dehydrogenase complex (PDC) in fatty acid biosynthesis supplies chloroplastic NADH (Fig. 1). In the cytosol, glucose-6-phosphate dehydrogenase (G6PD) and 6-phosphogluconate dehydrogenase (6PGD) in the oxidative pentose phosphate pathway (OPPP), non-phosphorylating glyceraldehyde-3-phosphate dehydrogenase (GAPN) in glycolysis, and isocitrate dehydrogenase (IDH) in amino acid biosynthesis supply NADPH while phosphorylating glyceraldehyde-3-phosphate dehydrogenase (GAPC), phosphoglycerate kinase (PGK), and pyruvate kinase (PK) in glycolysis supply NADH and ATP. In mitochondria, the photorespiratory glycine decarboxylase complex (GDC) and several enzymes of the tricarboxylic acid cycle including PDC, and IDH supply NADH.

**Figure 1.**
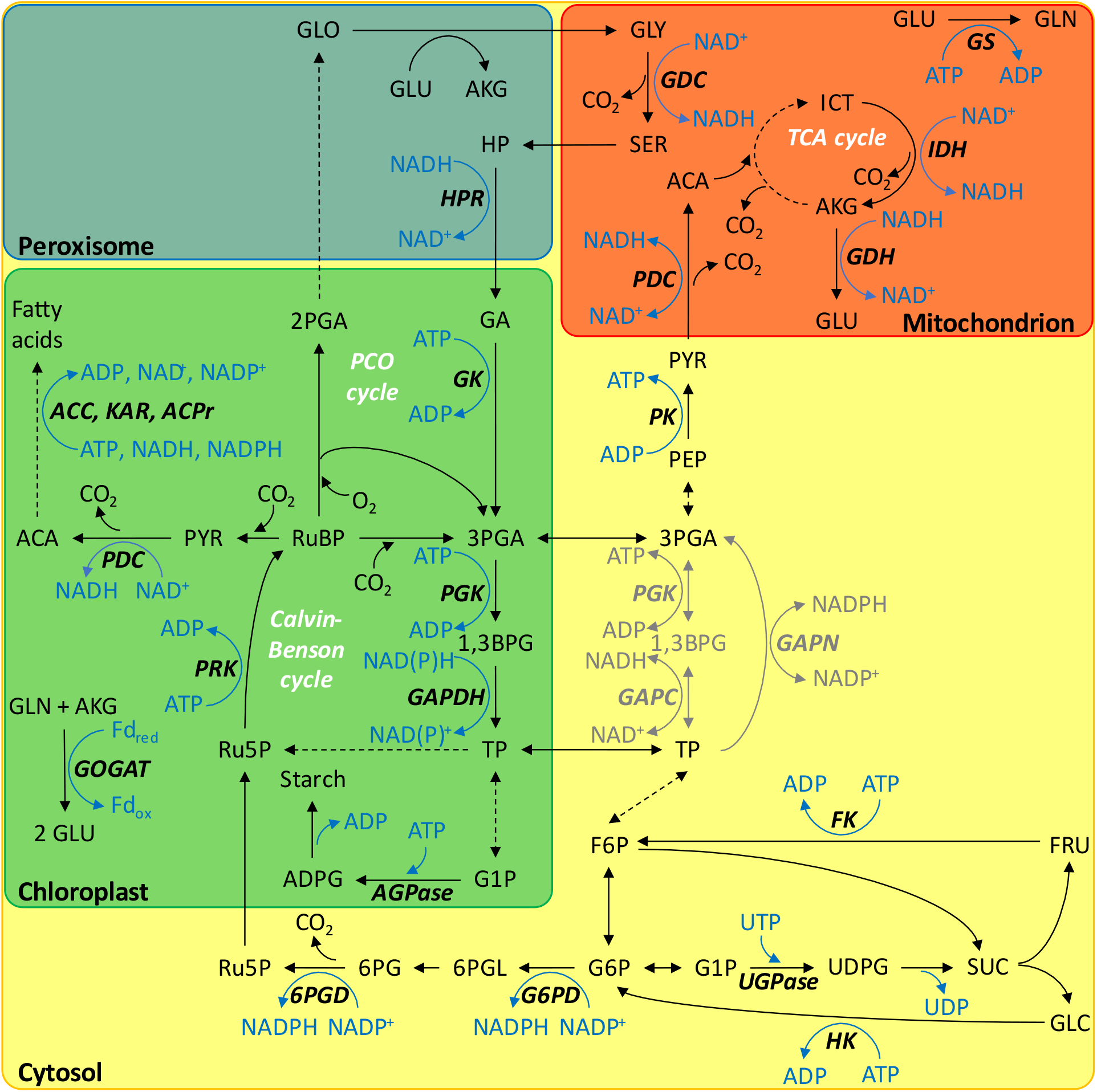
Sites of cofactor consumption and production in central carbon metabolism. Previously, we estimated carbon fluxes in 4-week-old photosynthesising *Camelina sativa* leaves (Xu et al. 2022). According to this analysis, pathways in black carry significant flux. Pathways in grey can be expected to carry significant flux but were not considered in our previous analysis. Enzymes: 6PGD, 6-phosphogluconate dehydrogenase; ACC, acetyl-CoA carboxylase; ACPr, 2,3-trans-enoyl-ACP reductase; AGPase, ADP-glucose pyrophosphorylase; FK, fructokinase; G6PD, glucose-6-phosphate dehydrogenase; GAPC (cytosolic) and GAPDH (chloroplastic), phosphorylating glyceraldehyde-3-phosphate dehydrogenase; GAPN, non-phosphorylating glyceraldehyde-3-phosphate dehydrogenase; GDC, glycine decarboxylase complex; GDH, glutamate dehydrogenase; GK, glycerate kinase; GOGAT, glutamine α-ketoglutarate aminotransferase; GS, glutamine synthetase; HK, hexokinase; HPR, hydroxypyruvate reductase; IDH, isocitrate dehydrogenase; KAR, 3-ketoacyl-ACP reductase; PDC, pyruvate dehydrogenase complex; PGK, phosphoglycerate kinase; PK, pyruvate kinase; PRK, phosphoribulokinase; UGPase, UDP-glucose pyrophosphorylase. Metabolites: 1,3BPG, 1,3-bisphosphoglycerate; 2PGA, 2-phosphoglycerate; 3PGA, 3-phosphoglycerate; 6PG, 6-phosphogluconate; 6PGL, 6-phosphogluconolactone; ACA, acetyl coenzyme A; ADPG, ADP-glucose; AKG, α-ketoglutarate; F6P, fructose 6-phosphate; FRU, fructose; G1P, glucose 1-phosphate; G6P, glucose 6-phosphate; GA, glycerate; GLC, glucose; GLN, glutamine; GLO, glyoxylate; GLU, glutamate; GLY, glycine; HP, hydroxypyruvate; ICT, isocitrate; PEP, phospho*enol*pyruvate; PYR, pyruvate; Ru5P, ribulose 5-phosphate; RuBP, ribulose 1,5-bisphosphate; SER, serine; SUC, sucrose; TP; triose phosphate (glyceraldehyde 3-phosphate and dihydroxyacetone phosphate); UDPG, UDP-glucose.

On the other hand, numerous enzyme reactions spread across several cell compartments consume reductant and energy cofactors. In chloroplasts, PGK, phosphorylating glyceraldehyde-3-phosphate dehydrogenase (GAPDH), and phosphoribulokinase (PRK) in the Calvin-Benson cycle (CBC) consume NAD(P)H, and ATP. Furthermore, glutamine α-ketoglutarate aminotransferase (GOGAT), and glycerate kinase (GK) in photorespiration, ADP-glucose pyrophosphorylase (AGPase) in starch biosynthesis, and acetyl-CoA carboxylase (ACC), 3-ketoacyl-ACP reductase (KAR), and 2,3-trans-enoyl-ACP reductase (ACPr) in fatty acid biosynthesis consume chloroplastic NAD(P)H, Fd_red_, and ATP. In the cytosol, UDP-glucose pyrophosphorylase (UGPase) in sucrose biosynthesis, and hexokinase (HK), and fructokinase (FK) in sucrose cycling consume UTP, and ATP. In mitochondria, glutamine synthetase (GS) in photorespiration, and glutamate dehydrogenase (GDH) in amino acid biosynthesis consume ATP, and NADH while photorespiratory hydroxypyruvate reductase (HPR) consumes NADH in peroxisomes.

Several mechanisms enable transmembrane transport of cofactors from sites of production to sites of consumption (Fig. 2). NAD^+^ carrier proteins enable direct transport of NAD(P)H across the inner membranes of chloroplasts and mitochondria but have low affinities for these reductants (Palmieri et al. 2009). By contrast, dicarboxylate transporters coupled with malate dehydrogenase enzymes (malate valves) enable indirect exchange of NAD(P)H among chloroplasts, peroxisomes, mitochondria, and the cytosol (Selinski and Scheibe 2019). While there is an NADPH-dependent malate dehydrogenase inside chloroplasts, most malate dehydrogenase provide NADH, not NADPH. By contrast, glyceraldehyde 3-phosphate/3-phosphoglycerate (GAP/3PGA) cycling enables NADPH transfer from chloroplasts to the cytosol (Kelly and Gibbs 1973b, a). In this cycle, chloroplastic PGK and GAPDH reduce 3PGA to GAP consuming ATP and NADPH. Subsequently, GAP is exported to the cytosol where GAPN is reducing it to 3PGA producing NADPH. A similar cycle involving GAPC, and PK (instead of GAPN) can provide cytosolic NADH and ATP at equimolar amounts (Stocking and Larson 1969). Additional cytosolic ATP can come from oxidative phosphorylation through the ADP/ATP carrier (Klingenberg 2008).

**Figure 2.**
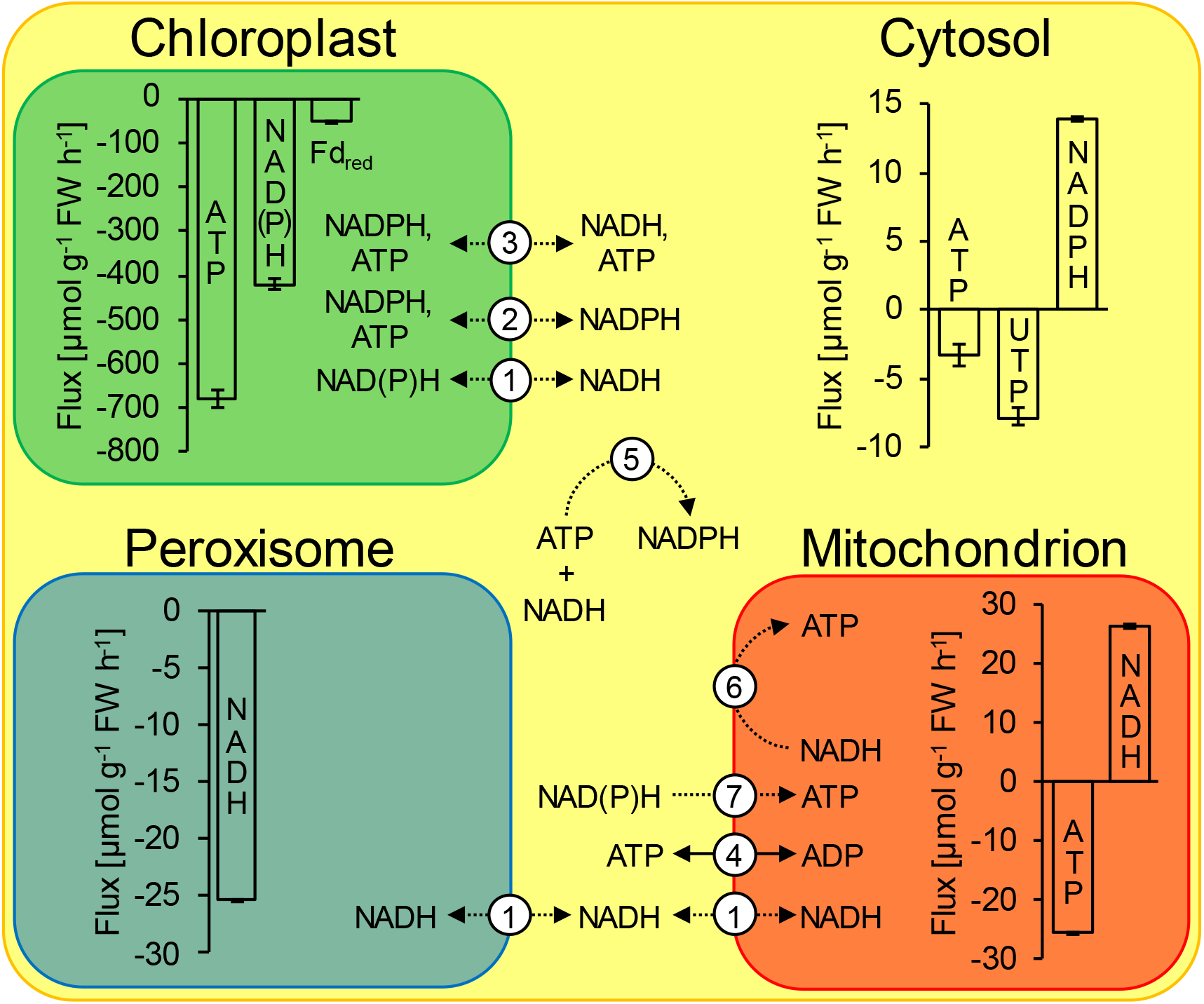
Compartment-specific cofactor consumption (negative values) and production (positive values) by central carbon metabolism in 4-week-old photosynthesising *Camelina sativa* leaves. The figure shows the net effect of all the processes listed in Table 2. Error bars represent 95% confidence intervals. 1, malate valves exchange NADH through malate-oxaloacetate interconversion by malate dehydrogenase (Selinski and Scheibe 2019). A chloroplastic isoform accepts NADPH. 2 and 3, cofactor transfer through redox cycles proposed by Kelly and Gibbs (1973a) and Stocking and Larson (1969), respectively. 4, counter-exchange of cytosolic ADP for mitochondrial ATP by the ADP/ATP carrier (Klingenberg 2008). 5, conversion of ATP and NADH to NADPH by the cytosolic oxidation-reduction cycle (Wieloch 2021). 6, NADH to ATP conversion by oxidative phosphorylation. 7, conversion of cytosolic NAD(P)H to mitochondrial ATP by oxidative phosphorylation starting at type II NAD(P)H dehydrogenases and glycerol-3-phosphate dehydrogenase both located in the inner mitochondrial membrane facing the cytosol. Solid arrow, direct cofactor transport. Dotted arrow, indirect cofactor transport via redox reactions.

Furthermore, several mechanisms enable cofactor conversion. In the cytosol, equimolar amounts of ATP, and NADH can be converted to NADPH by the carbon-neutral cytosolic oxidation-reduction (COR) cycle (Figs. 1–2; Wieloch 2021). In its forward direction, cytosolic GAPN oxidises GAP to 3PGA producing NADPH. In its backward direction, cytosolic PGK and GAPC reduce 3PGA to GAP consuming ATP, and NADH. COR cycling may help balancing the cytosolic NADPH supply according to varying demands. For instance, we recently reported evidence for increased COR cycling under drought where increased synthesis of reactive oxygen species may require increased amounts of cytosolic NADPH to maintain the cellular redox balance (Wieloch et al. 2021). In mitochondria, NADH can be converted to ATP by oxidative phosphorylation. Similarly, cytosolic NAD(P)H can be converted to mitochondrial ATP by oxidative phosphorylation starting not at complex I but at type II NAD(P)H dehydrogenases or glycerol-3-phosphate dehydrogenase both located in the inner mitochondrial membrane facing the cytosol (Rasmusson et al. 2008).

In a recently published study, we estimated carbon fluxes across central metabolism in illuminated *Camelina sativa* leaves, including flux through the CBC, photorespiration, glycolysis, the tricarboxylic acid cycle, and starch, sucrose, fatty acid, and amino acid biosynthesis (Xu et al. 2022). Importantly, our analyses explained the ^13^C labelling lag of CBC metabolites (a longstanding conundrum in plant biochemistry) by reinjection of weakly labelled carbon from vacuolar and cytosolic sugar pools into the CBC. This process involves breakdown of sucrose into fructose and glucose, ATP-dependent re-phosphorylation to fructose 6-phosphate and glucose 6-phosphate, UTP-dependent sucrose resynthesis from fructose 6-phosphate and glucose 6-phosphate and reinjection of carbon from cytosolic glucose 6-phosphate into the CBC via the NADPH-producing OPPP (Fig. 1). Additionally, assimilation of the CO_2_ released by the OPPP involves all cofactor-dependent reactions in the CBC and photorespiration (ATP, NAD(P)H, Fd_red_).

Here, we comprehensively characterise energy metabolism in illuminated *Camelina sativa* leaves based on the carbon flux estimates of our previous study (Xu et al. 2022). More specifically, we first estimate compartment-specific cofactor demands of sucrose cycling and the OPPP and discuss associated effects on plant carbon and energy metabolism. Subsequently, we estimate compartment-specific cofactor demands of central carbon metabolism and address the following questions (with respect to the conditions and set of reactions studied here):

- Is cofactor export from chloroplasts strictly required?
- Are mitochondria energetically self-sufficient?
- Does photorespiration have the same chloroplastic cofactor requirements as the CBC?
- How much cyclic electron flux is required to meet the cofactor demands of carbon metabolism?
- Where is the cytosolic ATP coming from?
- Is oxidative phosphorylation strictly required? (Note that oxidative phosphorylation does not strictly require tricarboxylic acid cycling.)
- Is mitochondrial NADH strictly required to balance peroxisomal NADH demands?
- Where is the cytosolic NADPH coming from?

## Material and methods

### Data

Previously, we estimated carbon fluxes across central metabolism in 4-week-old source leaves of wild-type *Camelina sativa* ecotype Suneson by ^13^C isotopically non-stationary metabolic flux analysis (Xu et al. 2022). These plants were grown at 22 °C, 50% relative humidity, and a light intensity of 500 μmol photons m^-2^ s^-1^ (8/16 h, day/night). Here, we utilise the carbon flux estimates to estimate associated cofactor fluxes (Fig. 1).

### Cofactor demand by carbon metabolism

We resolved chloroplastic (.p), cytosolic (.c), mitochondrial (.m), and peroxisomal (.ox) fluxes. Within the Calvin-Benson cycle, we considered reactions catalysed by PGK.p (−1 ATP per turnover), GAPDH.p (−1 NAD(P)H per turnover), and PRK.p (−1 ATP per turnover). Within photorespiration, we considered reactions catalysed by GS.m (−1 ATP per turnover), GOGAT.p (−2 Fd_red_ per turnover), GDC.m (+1 NADH per turnover), HPR.ox (−1 NADH per turnover), and GK.p (−1 ATP per turnover). It is commonly believed that photorespiratory ammonium is recaptured by GS2. While this enzyme is located in both chloroplasts and mitochondria (Taira et al. 2004), we here only considered the latter isoform since it has been suggested to immediately recapture potentially harmful ammonium released by GDC.m (Buchanan et al. 2015). Furthermore, while there is NADPH- and Fd_red_-dependent GOGAT.p, only the latter is essential for photorespiration and, therefore, considered here (Somerville and Ogren 1980). Within the OPPP, we considered reactions catalysed by G6PD.c (+1 NADPH per turnover), and 6PGD.c (+1 NADPH per turnover). Within starch and sucrose biosynthesis and sucrose cycling, we considered reactions catalysed by AGPase.p (−1 ATP per turnover), UGPase.c (−1 UTP per turnover), HK.c (−1 ATP per turnover), and FK.c (−1 ATP per turnover). Within glycolysis, we considered the reaction catalysed by PK.c (+1 ATP per turnover). Within the tricarboxylic acid cycle and amino acid biosynthesis, we only considered reactions catalysed by PDC.m (+1 NADH per turnover), IDH.m (+1 NADH per turnover), and GDH.m (−1 NADH per turnover) because other cofactor-dependent reactions reportedly carry negligible flux (Xu et al. 2022). Within fatty acid biosynthesis, we considered reactions catalysed by PDC.p (+1 NADH per turnover), ACC.p (−1 ATP per turnover), KAR.p (−1 NADPH per turnover), and ACPr.p (−1 NADH per turnover). Our accounting makes three assumptions. First, all pyruvate entering fatty acid biosynthesis comes from the β-elimination of phosphate from a carbocation intermediate of ribulose 1,5-bisphosphate (RuBP) carboxylation by rubisco (Andrews and Kane 1991). At 25 °C, the ratio of rubisco pyruvate synthesis to carboxylation is 0.7% (Andrews and Kane 1991). Xu et al. (2022) reported carboxylation proceeds at a rate of 172.1 μmol RuBP g^-1^ FW h^-1^ corresponding to ≈1.2 μmol g^-1^ FW h^-1^ pyruvate synthesis. Thus, pyruvate provided by rubisco significantly exceeds pyruvate requirements by fatty acid biosynthesis (0.4 μmol pyruvate g^-1^ FW h^-1^). Second, since the major isoform of ACPr.p is NADH-dependent and a minor isoform works with both NADH and NADPH (Buchanan et al. 2015), our accounting only considers NADH as cofactor of ACPr.p. Third, only C_16_ fatty acid are being produced. Initiation of C_16_ fatty acid biosynthesis requires 2 molecules pyruvate, 1 ATP, and 1 NADPH, and yields 1 C_4_ fatty acid, and 1 NADH. Elongation to a C_16_ fatty acid requires another 6 molecules pyruvate, 6 ATP, and 6 NADPH. Overall, biosynthesis of a C_16_ fatty acid requires 8 molecules pyruvate, 7 ATP, 7 NADPH and yields 1 NADH.

The cytosolic OPPP forms a shunt bypassing much of the CBC and releases CO_2_ from metabolism. We have calculated rubisco carboxylation and oxygenation rates accounting for the fixation of this CO_2_ as

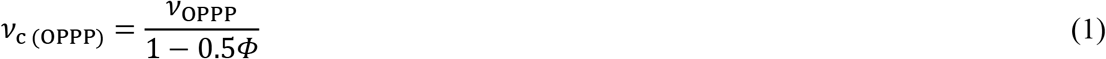

and

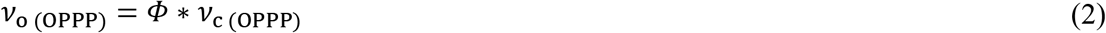

where *v*_OPPP_ denotes flux through the cytosolic OPPP, *v_c_* and *v_o_* denote rubisco carboxylation and oxygenation rates, respectively, *Φ* denotes the *v*_o_ / *v*_c_ ratio, and *v*_*c*(OPPP)_ and *v*_o(OPPP)_ denote rubisco carboxylation and oxygenation rates attributable to the CO_2_ lost in the OPPP, respectively. Please find the derivation of equation 1 in the supplement.

## Results and discussion

### Impacts of the cytosolic OPPP on plant carbon and energy metabolism

Flux through the cytosolic OPPP is likely a general feature of C3 metabolism since it can explain labelling lags of CBC metabolites (Xu et al. 2022) observed in several species fed ^13^CO_2_ or ^14^CO_2_ (Mahon et al. 1974; Canvin 1979; Hasunuma et al. 2010; Szecowka et al. 2013; Ma et al. 2014; Sharkey et al. 2020; Xu et al. 2021). In illuminated *Camelina sativa* leaves, the cytosolic OPPP supplies 14 μmol NADPH g^-1^ FW h^-1^, corresponding to 3.3% of the NAD(P)H consumed in the CBC (Tables 1–2). NADPH supply proceeds at about 10% of the rate of net carbon assimilation. This NADPH may support cytosolic redox, signalling, and biosynthetic metabolism and mitochondrial ATP production (Rasmusson et al. 2008).

**Table 1.**
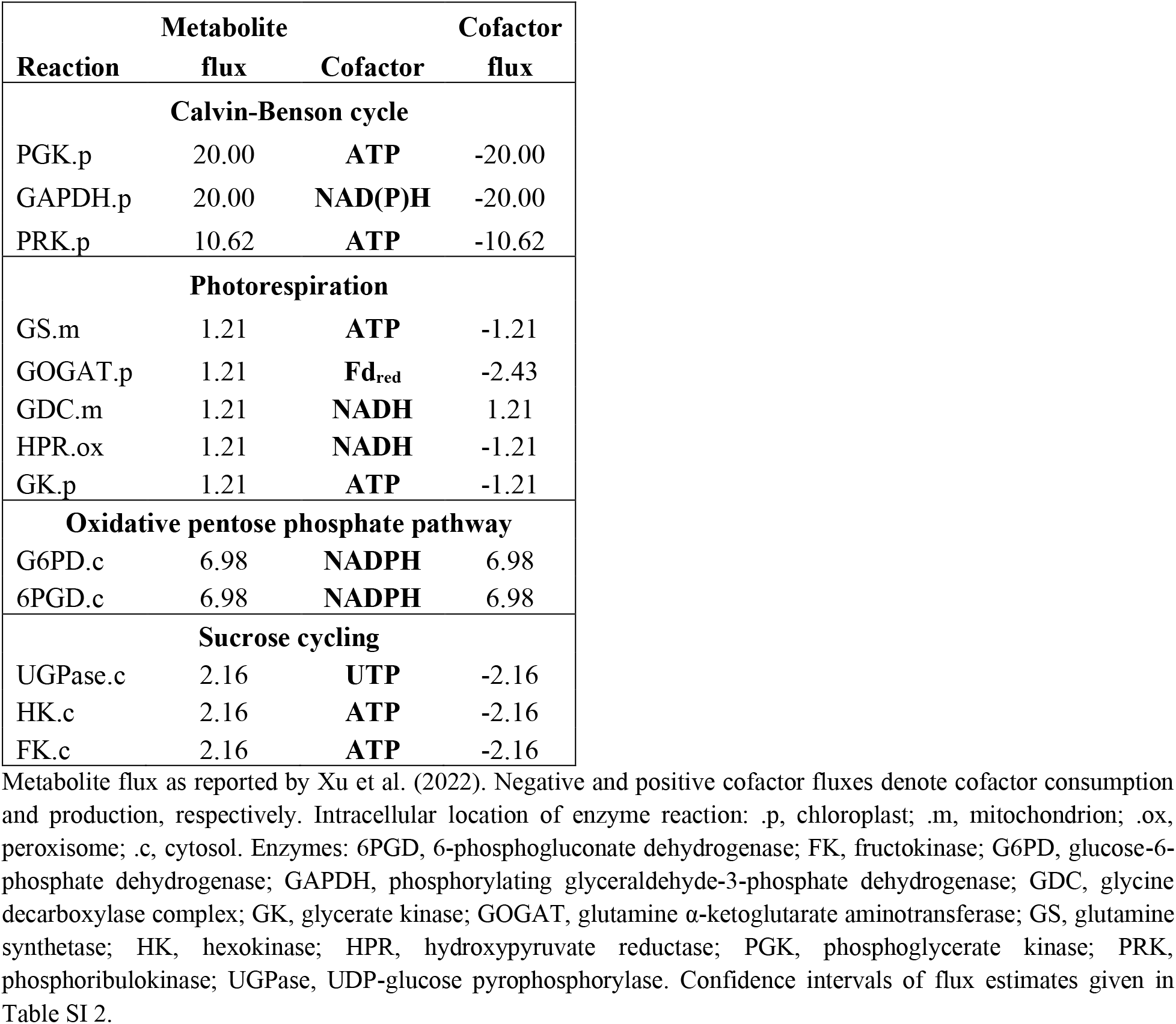
Metabolite and cofactor flux associated with sucrose cycling and carbon re-injection into the Calvin-Benson cycle by the cytosolic oxidative pentose phosphate pathway in illuminated *Camelina sativa* leaves [μmol g^-1^ FW h^-1^].

**Table 2.** Metabolite and cofactor flux in illuminated *Camelina sativa* leaves [μmol g^-1^ FW h^-1^].

| Reaction | Metabolite flux | Cofactor | Cofactor flux |
| --- | --- | --- | --- |
| **Calvin-Benson cycle** |  |  |  |
| PGK.p | 420.20 | ATP | -420.20 |
| GAPDH.p | 420.20 | NAD(P)H | -420.20 |
| PRK.p | 223.10 | ATP | -223.10 |
| **Photorespiration** |  |  |  |
| GS.m | 25.50 | ATP | -25.50 |
| GOGAT.p | 25.50 | Fd<sub>red</sub> | -51.00 |
| GDC.m | 25.49 | NADH | 25.49 |
| HPR.ox | 25.38 | NADH | -25.38 |
| GK.p | 25.38 | ATP | -25.38 |
| **Oxidative pentose phosphate pathway** |  |  |  |
| G6PD.c | 6.98 | NADPH | 6.98 |
| 6PGD.c | 6.98 | NADPH | 6.98 |
| **Starch and sucrose biosynthesis, and sucrose cycling** |  |  |  |
| AGPase.p | 10.51 | ATP | -10.51 |
| UGPase.c | 7.86 | UTP | -7.86 |
| HK.c | 2.16 | ATP | -2.16 |
| FK.c | 2.16 | ATP | -2.16 |
| **Glycolysis** |  |  |  |
| PK.c | 0.99 | ATP | 0.99 |
| **Fatty acid biosynthesis** |  |  |  |
| PDC.p | 0.44 | NADH | 0.44 |
| ACC.p | 0.44 | ATP | -0.39 |
| KAR.p | 0.44 | NADPH | -0.39 |
| ACPr.p | 0.44 | NADH | -0.39 |
| **Tricarboxylic acid cycle** |  |  |  |
| PDC.m | 0.92 | NADH | 0.92 |
| IDH.m | 0.92 | NADH | 0.92 |
| **Amino acid biosynthesis** |  |  |  |
| GDH.m | 0.92 | NADH | -0.92 |

Metabolite flux as reported by Xu et al. (2022). Negative and positive cofactor fluxes denote cofactor consumption and production, respectively. Intracellular location of enzyme reaction: .p, chloroplast; .m, mitochondrion; .ox, peroxisome; .c, cytosol. Enzymes: 6PGD, 6-phosphogluconate dehydrogenase; ACC, acetyl-CoA carboxylase; ACPr, 2,3-trans-enoyl-ACP reductase; AGPase, ADP-glucose pyrophosphorylase; FK, fructokinase; G6PD, glucose-6-phosphate dehydrogenase; GAPDH, phosphorylating glyceraldehyde-3-phosphate dehydrogenase; GDC, glycine decarboxylase complex; GDH, glutamate dehydrogenase; GK, glycerate kinase; GOGAT, glutamine α-ketoglutarate aminotransferase; GS, glutamine synthetase; HK, hexokinase; HPR, hydroxypyruvate reductase; IDH, isocitrate dehydrogenase; KAR, 3-ketoacyl-ACP reductase; PDC, pyruvate dehydrogenase complex; PGK, phosphoglycerate kinase; PK, pyruvate kinase; PRK, phosphoribulokinase; UGPase, UDP-glucose pyrophosphorylase. Confidence intervals of flux estimates given in Table SI 3.

Cytosolic NADPH production by the OPPP is associated with the release of 7 μmol CO_2_ g^-1^ FW h^-1^ from metabolism (Table S1). Fixation of this CO_2_ accounts for 4.8% of the total activity of rubisco. That is, 4.8% of the total flux through the CBC and photorespiration is spent on providing cytosolic NADPH, a significant investment. Associated energy requirements (20 μmol NADPH g^-1^ FW h^-1^, 2.4 μmol Fd_red_ g^-1^ FW h^-1^, 33 μmol ATP g^-1^ FW h^-1^) exceed the energy output of the OPPP (14 μmol NADPH g^-1^ FW h^-1^; Table 1). These carbon and energy costs would increase with increasing photorespiration. Thus, carbon metabolism in illuminated autotrophic tissue is not simply optimised for flux into carbon sinks but sacrifices both carbon and energy use efficiency to support cytosolic energy metabolism.

To try to reduce costs related to OPPP respiration, we suggest bioengineering and testing plants with a repressed cytosolic OPPP, and a newly introduced cytosolic NADPH-dependent malate dehydrogenase finely tuned to compensate for the loss in OPPP activity. This latter addition will only be required under conditions where carbon-neutral GAP/3PGA cycles and the COR cycle fall short of compensating for the reduction in NADPH from the OPPP.

### Impact of sucrose cycling on plant energy metabolism

Sucrose cycling involving breakdown of sucrose into fructose and glucose, ATP-dependent re-phosphorylation to fructose 6-phosphate and glucose 6-phosphate, and UTP-dependent sucrose resynthesis from fructose 6-phosphate and glucose 6-phosphate is another futile carbon cycle (Fig. 1). In the cytosol of illuminated *Camelina sativa* leaves, its operation requires 6.5 μmol ATP g^-1^ FW h^-1^ (Table 1; phosphorylation of 1 UDP to UTP consumes 1 ATP). This corresponds to 1% and 53% of the ATP requirement in chloroplasts and the cytosol, respectively (Table 2). Thus, sucrose cycling is a minor investment in overall leaf energy metabolism but a significant investment in cytosolic energy metabolism. However, it is important to note that several processes consuming cytosolic ATP are not considered here including cytosol-vacuole proton pumping (Hedrich et al. 2015). Therefore, it is unlikely that half the ATP demand in the cytosol comes from sucrose cycling.

### Cofactor fluxes in central carbon metabolism

Table 2 lists cofactor consumption and production by central enzyme reactions in carbon metabolism of *Camelina sativa* leaves. Figure 2 summarises this information for specific cell compartments. Chloroplast metabolism consumes large amounts of ATP and NADPH and some Fd_red_ while producing small amounts of NADH. Cytosol metabolism consumes ATP and UTP while producing NADPH. If triose phosphates were the only cytosolic carbon substrates (i.e., if 3PGA was not exported) then there would be an additional supply of 2.15 (+0.19, −0.16) μmol g^-1^ FW h^-1^ of NADH and ATP by GAPC and PGK or NADPH by GAPN (Fig. 1, grey reactions). Conversely, if exported 3PGA was reduced to triose phosphates, there would be an additional demand for cytosolic ATP and NADH by PGK and GAPC. However, these theoretical scenarios are not considered here since we have no estimates of carbon fluxes between cytosolic triose phosphates and 3PGA. Peroxisome metabolism consumes NADH, and mitochondrion metabolism consumes ATP while producing NADH. Compared to other cell compartments, absolute energy cofactor fluxes (ATP, UTP) in chloroplast are 26.7-times higher while absolute reductant cofactor fluxes (NAD(P)H, Fd_red_) are 17.9-times higher.

### Is cofactor export from chloroplasts strictly required?

Cofactor export from chloroplasts is mediated by the malate valve and GAP/3PGA cycles (Fig. 2). To investigate whether these processes are strictly required, we will now illustrate how cofactors produced by extra-chloroplastic carbon metabolism are used most efficiently to meet extra-chloroplastic cofactor demands. In this theoretical scenario, the mitochondrial NADH surplus (26.4 μmol g^-1^ FW h^-1^) is used to balance the peroxisomal NADH deficit (25.4 μmol g^-1^ FW h^-1^) through NADH transmembrane transport by malate valves (Fig. 2). Assuming a P/O ratio of 2.5 (Hinkle 2005), remaining mitochondrial NADH (1.03 μmol g^-1^ FW h^-1^) is converted to 2.6 μmol ATP g^-1^ FW h^-1^ via oxidative phosphorylation starting at complex I of the electron transport chain. Remaining mitochondrial and cytosolic ATP and UTP deficits equivalent to 34.1 μmol ATP g^-1^ FW h^-1^ are met by transmitting electrons from cytosolic NADPH onto the mitochondrial electron transport chain via type II NAD(P)H dehydrogenases. This reduces ATP yields by about 30% compared to transmission via complex I (Rasmusson et al. 2008). That is, the cytosolic surplus of 14 μmol NADPH g^-1^ FW h^-1^ is converted to 24.4 μmol ATP g^-1^ FW h^-1^ in mitochondria which is partly exported to the cytosol via the ADP/ATP carrier (Klingenberg 2008). Overall, there remains an extra-chloroplastic deficit of 9.7 μmol ATP g^-1^ FW h^-1^. Thus, with respect to the conditions and set of reactions studied here, cofactor export from chloroplasts is strictly required to meet all extra-chloroplastic cofactor demands.

### Are mitochondria energetically self-sufficient?

It may be most realistic to assume that the mitochondrial surplus of 26.4 μmol NADH g^-1^ FW h^-1^ is used first to balance the mitochondrial deficit of 25.5 μmol ATP g^-1^ FW h^-1^ rather than being directly shuttled out of mitochondria into peroxisomes (Fig. 2). To meet the mitochondrial ATP deficit by oxidative phosphorylation starting from complex I requires only 10.2 μmol NADH g^-1^ FW h^-1^. Thus, with respect to the conditions and set of reactions studied here, mitochondria are energetically self-sufficient.

### Does photorespiration have the same chloroplastic cofactor requirements as the CBC?

Our cofactor accounting assumes that photorespiratory ammonium is recaptured by mitochondrial (not chloroplastic) GS. This has implications for the cofactor requirements in chloroplasts. Chloroplastic cofactor requirements upon carboxylation of 2 RuBP comprise 4 ATP by PGK, 4 NADPH by GAPDH, and 2 ATP by PRK. Chloroplastic cofactor requirements upon oxygenation of 2 RuBP comprise 3 ATP by PGK, 3 NADPH by GAPDH, 2 ATP by PRK, 1 ATP by GK, and 2 Fd_red_ corresponding to 1 NADPH by GOGAT. Thus, overall, both processes have the same absolute chloroplastic cofactor requirements with a chloroplastic ATP/NADPH demand ratio of 1.5. Other processes, such as starch biosynthesis and serine withdrawal from photorespiration, will cause only small changes of this ratio (see below).

### How much cyclic electron flux is required to meet the cofactor demands of carbon metabolism?

Chloroplast carbon metabolism has much higher demands for reductant and energy cofactors than any other cell compartment (Fig. 2). These demands are readily satisfied by cofactor input mainly from the light reactions of photosynthesis without the need for transmembrane transport or conversion of cofactors. Interestingly, chloroplast GAPDH accepts both NADPH and NADH (McGowan and Gibbs 1974). *In vivo*, however, NADPH supply from the light reactions is 7630-times higher than the NADH surplus from fatty acid biosynthesis (Table 2).

Overall, chloroplast metabolism requires 680 μmol ATP g^-1^ FW h^-1^ and 446 μmol NAD(P)H g^-1^ FW h^-1^ (Table 2). This accounts for Fd_red_ demands where 2 Fd_red_ correspond to 1 NADPH. Hence, the reaction network and conditions studied here have an ATP/NADPH demand ratio of 1.52. However, linear electron flux including Q cycling has an estimated ATP/NADPH supply ratio of 1.286 (assuming trans-membrane transport of 3 protons per electron and production of 3 ATP per 14 protons by ATP synthase) (Seelert et al. 2000). Thus, assuming neither the water-water cycle (affecting the ATP/NADPH supply ratio) nor GAP/3PGA cycles and the chloroplast malate valve (affecting the ATP/NADPH demand ratio) are operational, cyclic electron flux through photosystem I is required.

Production of 446 μmol NADPH g^-1^ FW h^-1^ requires linear electron flux at a rate of 892 μmol g^-1^ FW h^-1^ and yields 574 μmol ATP g^-1^ FW h^-1^. To meet the remaining deficit of 106 μmol ATP g^-1^ FW h^-1^, cyclic electron flux through photosystem I including Q cycling at a rate of 248 μmol g^-1^ FW h^-1^ is required (assuming trans-membrane transport of 2 protons per electron and production of 3 ATP per 14 protons by ATP synthase) (Seelert et al. 2000). Thus, with respect to the conditions and set of reactions studied here, cyclic electron flux accounts for 21.7% of whole electron flux and is a 27.8% fraction of linear electron flux.

Jointly, linear, and cyclic electron flux through photosystem I consume 2032 μmol photons g^-1^ FW h^-1^. Carbon flux in *Camelina sativa* leaves was measured at 500 μmol photons m^-2^ s^-1^ corresponding to 3273 μmol photons g^-1^ FW h^-1^ (the fresh weight per leaf area for a typical Camelina leaf is 0.055 ± 0.0041 g cm^-2^ implying a leaf thickness of 0.55 mm). Thus, with respect to the conditions and set of reactions studied here, 62% of the photons falling on the leaf are absorbed and used for photosynthesis.

In our previous analysis (Xu et al. 2022), carbon fluxes through GAP/3PGA cycles, the COR cycle (grey in Fig. 1), and the chloroplast malate valve were not estimated, i.e., associated cofactor fluxes are unknown. These processes would, however, decrease the ATP/NADPH demand ratio of carbon metabolism and, thus, the requirement for cyclic electron flux. This is because GAP/3PGA cycles require chloroplastic ATP and NAD(P)H at a 1:1 ratio (Stocking and Larson 1969; Kelly and Gibbs 1973a), and the malate valve requires chloroplastic NAD(P)H only (Selinski and Scheibe 2019). COR cycling requires NADH and ATP in the cytosol which may be provided by combined action of the chloroplast malate valve and oxidative phosphorylation (Wieloch 2021). Furthermore, these processes would consume additional electrons and, thus, increase the photon use efficiency of electron transport. For example, no cyclic electron flux is required if 82.5 μmol NAD(P)H g^-1^ FW h^-1^ (corresponds to ≈60% of net CO_2_ assimilation) are exported by the chloroplast malate valve. This additional flux would result in 64.6% photon use efficiency of electron transport.

### Where is the cytosolic ATP coming from? Is oxidative phosphorylation strictly required?

Overall, we estimated an extra-chloroplastic ATP (includes UTP) deficit of 36.7 μmol g^-1^ FW h^-1^ (Fig. 2, Table 2) which may be balanced by the following mechanisms. First, cytosolic NADPH from the OPPP (14 μmol g^-1^ FW h^-1^) may be converted to 24.4 μmol ATP g^-1^ FW h^-1^ via oxidative phosphorylation starting at type II NADPH dehydrogenase (yields 2.5*0.7 ATP per NADPH). This leaves an ATP deficit of 12.3 μmol g^-1^ FW h^-1^. Second, GAP/3PGA cycling involving cytosolic GAPN may supply cytosolic NADPH (Fig. 1). To balance the entire ATP deficit of 36.7 μmol g^-1^ FW h^-1^ via oxidative phosphorylation starting at type II NADPH dehydrogenases would require a flux of 21 μmol GAP g^-1^ FW h^-1^ through this pathway. Third, mitochondrial NADH from photorespiration (26.4 μmol g^-1^ FW h^-1^) may be converted to up to 66 μmol ATP g^-1^ FW h^-1^ via oxidative phosphorylation starting at complex I of the mitochondrial electron transport chain (yields 2.5 ATP per NADH). Fourth, GAP/3PGA cycling involving cytosolic PGK may supply ATP (Fig. 1). This pathway provides ATP and NADH (from GAPC) at equimolar amounts. If the latter is imported into mitochondria by the malate valve and used for oxidative phosphorylation (Fig. 2), GAP/3PGA cycling at a rate of 10.5 μmol g^-1^ FW h^-1^ would suffice to meet the ATP deficit of 36.7 μmol g^-1^ FW h^-1^ (1 ATP from PGK, 2.5 ATP from NADH provided by GAPC). If NADH from GAPC enters oxidative phosphorylation via type II NADH dehydrogenase which reduces ATP yields by about 30% (see above), GAP/3PGA cycling at a rate of 13.3 μmol g^-1^ FW h^-1^ is required to meet the ATP deficit (1 ATP from PGK, 2.5*0.7 ATP from NADH). Fifth, GAP/3PGA cycling involving cytosolic PGK may supply all extra-chloroplastic ATP required (Fig. 1). This would provide an additional 36.7 μmol NADH g^-1^ FW h^-1^ from GAPC which could be used to balance the peroxisomal NADH deficit (25.4 μmol g^-1^ FW h^-1^). Remaining extra-chloroplastic NADH (37.7 μmol g^-1^ FW h^-1^, includes the mitochondrial NADH surplus) would then most likely enter oxidative phosphorylation. Sixth, cytosolic NADH supplied by the chloroplast malate valve may be converted to ATP by mitochondrial NADH import (malate valve) and oxidative phosphorylation or oxidative phosphorylation via type II NADH dehydrogenase. Supply of 36.7 μmol ATP g^-1^ FW h^-1^ by these pathways requires net flux through the chloroplast malate valve of 14.7 or 21 μmol NADH g^-1^ FW h^-1^, respectively. Hence, numerous metabolic pathways can supply cytosolic ATP. All of them, however, involve oxidative phosphorylation. Thus, in illuminated *Camelina sativa* leaves under the conditions studied here, oxidative phosphorylation is strictly required. This is in agreement with results from a recent simulation study (Shameer et al. 2019).

### Is mitochondrial NADH strictly required to balance peroxisomal NADH demands?

We estimated a peroxisomal NADH deficit of 25.4 μmol g^-1^ FW h^-1^ due to photorespiration (Fig. 2, Table 2). This deficit is often assumed to be balanced by the mitochondrial NADH surplus of 26.4 μmol g^-1^ FW h^-1^ coming mostly from photorespiration. However, there are other processes that may help to balance the peroxisomal NADH deficit. First, NAD(P)H from chloroplasts may be transferred to peroxisomes via malate valves. Second, combined action of GAP/3PGA cycling involving cytosolic GAPC and the peroxisomal malate valve my balance peroxisomal NADH demands. Concomitantly supplied ATP by PGK may help to balance extra-chloroplastic ATP deficits. Thus, to balance peroxisomal NADH demands, mitochondrial NADH is not strictly required. This is in agreement with results from a recent simulation study (Shameer et al. 2019). However, peroxisomal NADH import appears to be strictly required.

### Where is the cytosolic NADPH coming from?

In the cytosol of illuminated *Camelina sativa* leaves, 14 μmol NADPH g^-1^ FW h^-1^ are supplied by the OPPP. Primary flux control through the OPPP is exerted at the level of its first enzyme, G6PD. In the cytosol of Arabidopsis leaves, only G6PD5 was found (Wakao and Benning 2005). Reportedly, G6PD5 activity is not directly affected by redox agents (Wakao and Benning 2005). Correspondingly, cytosolic G6PD activity in *Solanum tuberosum* leaf discs was found to be constant in response to increasing H_2_O_2_ (Hauschild and von Schaewen 2003). By contrast, increases in activity occur with increasing sugar availability (sugars tested: mannose, glucose, fructose, sucrose) through *de novo* enzyme synthesis (Hauschild and von Schaewen 2003). As carbon assimilation decreases, so does sugar availability (Lawlor and Fock 1977; Sánchez-Rodríguez et al. 1999). This can be expected to result in decreasing OPPP flux and associated NADPH supply. However, decreasing carbon assimilation (e.g., due to drought) also promotes the synthesis of reactive oxygen species (Noctor et al. 2002) and their detoxification can be expected to increase the demand for cytosolic NADPH. Thus, especially (but not exclusively) under low carbon assimilation, other sources of cytosolic NADPH may be required.

GAPN catalyses the irreversible oxidation of GAP to 3PGA and simultaneous reduction of NADP^+^ to NADPH (Fig. 1). Interestingly, a GAPN null mutant showed increased expression and activity of cytosolic G6PD5 (3.8-fold increase in At3g27300) and an increased oxidative load (Rius et al. 2006). This suggests that G6PD5 and GAPN interact in redox control and that GAPN is required in redox control since increased G6PD5 expression cannot fully compensate for the loss in GAPN function.

Just like GAPN, GAPC and PGK catalyse cytosolic GAP to 3PGA conversions. Intriguingly, the GAPN null mutant showed increased expression and activity of GAPC (GAPC1: 3.5-fold increase in At3g04120) (Rius et al. 2006). This was suggested to compensate for the lack of GAPN activity (Rius et al. 2006). However, GAPC and PGK supply NADH and ATP which (in contrast to NADPH) cannot be used in redox control. Alternatively, upregulation of GAPC may promote COR cycling (Wieloch 2021). Each turn of this cycle provides NADPH at GAPN and consumes ATP and NADH at PGK and GAPC (Fig. 1). Required NADH and ATP may be provided via the chloroplast malate valve and oxidative phosphorylation (Fig. 2). In this regard, a putative moonlighting function of GAPC is worth noting (Scheibe 2019). With increasing oxidative load (as observed in the GAPN null mutant), GAPC is increasingly deactivated and moved to the nucleus (Piattoni et al. 2013; Hildebrandt et al. 2015). There, it can bind to a gene coding for chloroplast malate dehydrogenase, and expression of this gene was found to increase during the initial stages of oxidative stress (Hildebrandt et al. 2015). Furthermore, it is worth noting that the GAPN null mutant shows increased expression of cytosolic malate dehydrogenase (Rius et al. 2006). Taken together, this may result in increased malate valve capacity to support GAPC and PGK in COR cycling.

The most marked difference between cytosolic NADPH supply by the OPPP versus COR cycling is that the latter does not release CO_2_. Thus, cytosolic NADPH supply by COR cycling might be physiologically beneficial when CO_2_ assimilation is impeded, e.g., under photorespiratory conditions. In line with this reasoning, we recently reported isotope evidence for upregulation of COR cycling under drought (Wieloch et al. 2021). Under these same conditions, regulatory properties of G6PD5 suggest decreased flux through the OPPP (see above). Taken together, we propose that the OPPP and COR cycling act in concert to ensure the cytosolic NADPH supply under varying environmental/developmental conditions. To follow up on this proposal, wide-screening enzyme expression in a G6PD5 knockout mutant may be valuable.

## Author Contributions

T.W., and T.D.S. designed the research. T.W. led the research. T.W. analysed and interpreted the data and wrote the manuscript with input from T.D.S.

## Data Availability

All data generated or analysed during this study are included in this published article.

## Acknowledgments

T.D.S. was supported by the Division of Chemical Sciences, Geosciences and Biosciences, Office of Basic Energy Sciences of the United States Department of Energy Grant DE-FG02-91ER20021. T.D.S. receives partial salary support from MSU AgBioResearch. However, the contents of this paper are solely the responsibility of the authors.

## Notes

### Competing Interest Statement

The authors have declared no competing interest.

