## Supplemental information for "Compartment-specific energy requirements of photosynthetic carbon metabolism in *Camelina sativa* leaves"

### Derivation of equation 1

In the absence of carbon loss due to respiration, net carbon fixation via the Calvin-Benson cycle and photorespiration, $A$, is given as

$$A = v_{\mathrm{c}} - 0.5v_{\mathrm{o}} = V_{\mathrm{c}}(1 - 0.5\Phi) \quad \text{(S1)}$$

where $v_{\mathrm{c}}$ and $v_{\mathrm{o}}$ denote rubisco carboxylation and oxygenation rates, respectively, and $\Phi$ denotes the $v_{\mathrm{o}}$ / $v_{\mathrm{c}}$ ratio (Farquhar et al. 1980). Solving equation S1 for $v_{\mathrm{c}}$ yields

$$v_{\mathrm{c}} = \frac{A}{1 - 0.5\Phi} \quad \text{(S2)}$$

To calculate the rubisco carboxylation rate attributable to the $CO_2$ lost in the OPPP, $v_{\mathrm{c\ (OPPP)}}$, we substitute $v_{\mathrm{c}}$ for $v_{\mathrm{c\ (OPPP)}}$ and $A$ for the flux through the OPPP, $v_{\mathrm{OPPP}}$, which yields equation 1.

### Data tables

**Table S1** Flux and flux ratios related to rubisco and the cytosolic oxidative pentose phosphate pathway in illuminated *Camelina sativa* leaves.

| | **Flux or flux ratio** | **95% CI** | |
|---|---|---|---|
| | | **LB** | **UB** |
| $v_c$ | 172.10 | 166.43 | 178.75 |
| $v_o$ | 51.00 | 51.00 | 51.00 |
| $v_o / v_c = \Phi$ | 0.30 | 0.29 | 0.31 |
| $v_{OPPP}$ | 6.98 | 6.92 | 7.03 |
| $v_{c\,(OPPP)}$ | 8.19 | 8.07 | 8.30 |
| $v_{o\,(OPPP)}$ | 2.43 | 2.30 | 2.54 |
| $v_{c\,(OPPP)} / v_c$ | 0.048 | 0.045 | 0.050 |
| $v_{o\,(OPPP)} / v_o$ | 0.048 | 0.045 | 0.050 |

Flux is given in units of µmol $g^{-1}$ FW $h^{-1}$. Abbreviations: $v_c$, rubisco carboxylation rate; $v_o$, rubisco oxygenation rate; $\Phi$, $v_o / v_c$ ratio; $v_{OPPP}$, flux through the cytosolic oxidative pentose phosphate pathway; $v_{c\,(OPPP)}$ and $v_{o\,(OPPP)}$, $v_c$ and $v_o$ accounting for the net fixation of the $CO_2$ released by the cytosolic oxidative pentose phosphate pathway.

**Table S2** Metabolite and cofactor flux associated with sucrose cycling and carbon re-injection into the Calvin-Benson cycle by the cytosolic oxidative pentose phosphate pathway in illuminated *Camelina sativa* leaves [μmol $g^{-1}$ FW $h^{-1}$].

| Reaction | Metabolite flux | 95% CI LB | 95% CI UB | Cofactor | Cofactor flux | 95% CI LB | 95% CI UB |
|---|---|---|---|---|---|---|---|
| **Calvin-Benson cycle** | | | | | | | |
| PGK.p | 20.00 | 18.46 | 21.62 | **ATP** | -20.00 | -21.62 | -18.46 |
| GAPDH.p | 20.00 | 18.46 | 21.62 | **NAD(P)H** | -20.00 | -21.62 | -18.46 |
| PRK.p | 10.62 | 9.82 | 11.46 | **ATP** | -10.62 | -11.46 | -9.82 |
| **Photorespiration** | | | | | | | |
| GS.m | 1.21 | 1.15 | 1.27 | **ATP** | -1.21 | -1.27 | -1.15 |
| GOGAT.p | 1.21 | 1.15 | 1.27 | **$Fd_{red}$** | -2.43 | -2.54 | -2.30 |
| GDC.m | 1.21 | 1.15 | 1.27 | **NADH** | 1.21 | 1.15 | 1.27 |
| HPR.ox | 1.21 | 1.15 | 1.27 | **NADH** | -1.21 | -1.27 | -1.15 |
| GK.p | 1.21 | 1.15 | 1.27 | **ATP** | -1.21 | -1.27 | -1.15 |
| **Oxidative pentose phosphate pathway** | | | | | | | |
| G6PD.c | 6.98 | 6.92 | 7.03 | **NADPH** | 6.98 | 6.92 | 7.03 |
| 6PGD.c | 6.98 | 6.92 | 7.03 | **NADPH** | 6.98 | 6.92 | 7.03 |
| **Sucrose cycling** | | | | | | | |
| UGPase.c | 2.16 | 1.84 | 2.53 | **UTP** | -2.16 | -2.53 | -1.84 |
| HK.c | 2.16 | 1.84 | 2.53 | **ATP** | -2.16 | -2.53 | -1.84 |
| FK.c | 2.16 | 1.84 | 2.53 | **ATP** | -2.16 | -2.53 | -1.84 |

Metabolite flux as reported by Xu et al. (2022). Negative and positive cofactor fluxes denote cofactor consumption and production, respectively. Intracellular location of enzyme reaction: .p, chloroplast; .m, mitochondrion; .ox, peroxisome; .c, cytosol. Enzymes: 6PGD, 6-phosphogluconate dehydrogenase; FK, fructokinase; G6PD, glucose-6-phosphate dehydrogenase; GAPDH, phosphorylating glyceraldehyde-3-phosphate dehydrogenase; GDC, glycine decarboxylase complex; GK, glycerate kinase; GOGAT, glutamine α-ketoglutarate aminotransferase; GS, glutamine synthetase; HK, hexokinase; HPR, hydroxypyruvate reductase; PGK, phosphoglycerate kinase; PRK, phosphoribulokinase; UGPase, UDP-glucose pyrophosphorylase.

**Table S3** Metabolite and cofactor flux in illuminated *Camelina sativa* leaves [µmol $g^{-1}$ FW $h^{-1}$].

| Reaction | Metabolite flux | 95% CI LB | 95% CI UB | Cofactor | Cofactor flux | 95% CI LB | 95% CI UB |
|---|---|---|---|---|---|---|---|
| **Calvin-Benson cycle** | | | | | | | |
| PGK.p | 420.20 | 408.80 | 433.42 | **ATP** | -420.20 | -433.42 | -408.80 |
| GAPDH.p | 420.20 | 408.80 | 433.42 | **NAD(P)H** | -420.20 | -433.42 | -408.80 |
| PRK.p | 223.10 | 217.43 | 229.75 | **ATP** | -223.10 | -229.75 | -217.43 |
| **Photorespiration** | | | | | | | |
| GS.m | 25.50 | 25.50 | 25.50 | **ATP** | -25.50 | -25.50 | -25.50 |
| GOGAT.p | 25.50 | 25.50 | 25.50 | **$Fd_{red}$** | -51.00 | -51.00 | -51.00 |
| GDC.m | 25.49 | 25.49 | 25.49 | **NADH** | 25.49 | 25.49 | 25.49 |
| HPR.ox | 25.38 | 25.37 | 25.39 | **NADH** | -25.38 | -25.39 | -25.37 |
| GK.p | 25.38 | 25.37 | 25.39 | **ATP** | -25.38 | -25.39 | -25.37 |
| **Oxidative pentose phosphate pathway** | | | | | | | |
| G6PD.c | 6.98 | 6.92 | 7.03 | **NADPH** | 6.98 | 6.92 | 7.03 |
| 6PGD.c | 6.98 | 6.92 | 7.03 | **NADPH** | 6.98 | 6.92 | 7.03 |
| **Starch and sucrose biosynthesis, and sucrose cycling** | | | | | | | |
| AGPase.p | 10.51 | 10.51 | 10.51 | **ATP** | -10.51 | -10.51 | -10.51 |
| UGPase.c | 7.86 | 7.27 | 8.56 | **UTP** | -7.86 | -8.56 | -7.27 |
| HK.c | 2.16 | 1.84 | 2.53 | **ATP** | -2.16 | -2.53 | -1.84 |
| FK.c | 2.16 | 1.84 | 2.53 | **ATP** | -2.16 | -2.53 | -1.84 |
| **Glycolysis** | | | | | | | |
| PK.c | 0.99 | 0.91 | 1.08 | **ATP** | 0.99 | 0.91 | 1.08 |
| **Fatty acid biosynthesis** | | | | | | | |
| PDC.p | 0.44 | 0.44 | 0.44 | **NADH** | 0.44 | 0.44 | 0.44 |
| ACC.p | 0.44 | 0.44 | 0.44 | **ATP** | -0.39 | -0.39 | -0.39 |
| KAR.p | 0.44 | 0.44 | 0.44 | **NADPH** | -0.39 | -0.39 | -0.39 |
| ACPr.p | 0.44 | 0.44 | 0.44 | **NADH** | -0.39 | -0.39 | -0.39 |
| **Tricarboxylic acid cycle** | | | | | | | |
| PDC.m | 0.92 | 0.85 | 1.00 | **NADH** | 0.92 | 0.85 | 1.00 |
| IDH.m | 0.92 | 0.85 | 1.00 | **NADH** | 0.92 | 0.85 | 1.00 |
| **Amino acid biosynthesis** | | | | | | | |
| GDH.m | 0.92 | 0.85 | 1.00 | **NADH** | -0.92 | -1.00 | -0.85 |

Metabolite flux as reported by Xu et al. (2022). Negative and positive cofactor fluxes denote cofactor consumption and production, respectively. Intracellular location of enzyme reaction: .p, chloroplast; .m, mitochondrion; .ox, peroxisome; .c, cytosol. Enzymes: 6PGD, 6-phosphogluconate dehydrogenase; ACC, acetyl-CoA carboxylase; ACPr, 2,3-trans-enoyl-ACP reductase; AGPase, ADP-glucose pyrophosphorylase; FK, fructokinase; G6PD, glucose-6-phosphate dehydrogenase; GAPDH, phosphorylating glyceraldehyde-3-phosphate dehydrogenase; GDC, glycine decarboxylase complex; GDH, glutamate dehydrogenase; GK, glycerate kinase; GOGAT, glutamine α-ketoglutarate aminotransferase; GS, glutamine synthetase; HK, hexokinase; HPR, hydroxypyruvate reductase; IDH, isocitrate dehydrogenase; KAR, 3-ketoacyl-ACP reductase; PDC, pyruvate dehydrogenase complex; PGK, phosphoglycerate kinase; PK, pyruvate kinase; PRK, phosphoribulokinase; UGPase, UDP-glucose pyrophosphorylase.
